# Two-photon *in vivo* imaging reveals cell type-specific mitophagy dynamic changes in mouse somatosensory cortex during aging

**DOI:** 10.64898/2025.12.17.694842

**Authors:** Beatriz Escobar-Doncel, Xiaoyi Zhang, Mario Fernández de la Puebla, Shreyas Balachandra Rao, Synnøve Algrøy Fjeldstad, Maja Tvedt Dahle, Mahmood Amiry-Moghaddam, Magnar Bjørås, Evandro Fei Fang, Wannan Tang

**Author notes:** These authors contributed equally (Beatriz Escobar-Doncel, Xiaoyi Zhang). Correspondences (W.T.) and (E.F.F.). **Emails** Beatriz Escobar-Doncel; Xiaoyi Zhang; Mario Fernández de la Puebla; Shreyas Balachandra Rao; Synnøve Algrøy Fjeldstad; Maja Tvedt Dahle; Mahmood Amiry-Moghaddam; Magnar Bjørås; Evandro Fei Fang; Wannan Tang.

## Abstract

Mitochondrial homeostasis is maintained through mitochondrial autophagy, a process commonly known as mitophagy. Recent research has begun to highlight the region- and cell type-specific nature of mitophagy during brain aging; however, these dynamics have largely remained unexplored in living brains. To address this gap, we conducted two-photon mt-Keima imaging in somatosensory cortical neurons and astrocytes in behaving mice across two age groups. Our observations revealed a reduction in mitophagy levels in both cell types during aging, and we consistently found a higher level of mitophagy in astrocytes compared to neurons across both age groups, indicating a cell type-specific mitophagy dynamics that is independently of the aging process. NAD^+^ augmentation improved mitophagy level in both neurons and astrocytes in old-aged mice at the dose and method of administration tested. Collectively, our data underscore the critical importance of both neuronal and astrocytic metabolic roles in brain aging. The findings also suggest that NAD^+^ repletion has clinical value in potentially improving these cell type-specific mitophagy dynamics during aging.

## Introduction

Considering the brain as a highly energetic-demanding organ, growing research links dysfunctional mitochondria to the physical changes and neuronal degeneration observed in brain during aging.^1–3^ Mitochondria are essential organelles for maintaining cellular energy homeostasis, as well as other fundamental processes such as phospholipid biosynthesis/transport, calcium signaling, ion homeostasis, and even the arbitration of cell death or survival.^4–6^ In various model organisms, a reduction in mitochondrial function may contribute to the age-related decline in organ performance, which is a key factor in common neurodegenerative conditions like Alzheimer’s disease.^5,7^ Consequently, quality control mechanisms have evolved to restore and preserve energy metabolism, especially within the sensitive environment of the brain.^8,9^ During the selective removal of damaged mitochondria, termed mitophagy, a portion of the mitochondrial network is engulfed within double-membraned vesicles, which fuse with lysosomes for degradation of the engulfed cargo.^10^ The molecular mechanisms of mitophagy have also been elucidated in considerable detail in recent years.^11,12^

Various tools exist for measuring mitophagy, including electron microscopy,^13^ MitoTimer,^14^ co-localization of mitochondrial proteins and autophagic machinery,^15^ isolation of autophagic vesicles,^16,17^ and ratiometric fluorescent based probes, such as mito-Rosella,^18^ mitoQC,^19^ and mt-Keima.^20,21^ While these methods have distinct strengths and limitations, ratiometric probes are widely used for *ex vivo* assessments of mitophagic flux due to their high sensitivity and specificity. However, most measurements using these probes have been performed in fixed tissues.^18–20,22^ This approach sacrifices physiological context, as key regulatory signals (e.g., metabolism, temperature) are lost upon tissue extraction, preventing accurate capture of dynamic status of mitophagy regulation and compromising the physiological relevance of the findings.^23^ Furthermore, most of the *ex vivo* brain mitophagy measurements were either cell type unspecific or targeting only neurons, leaving other cell types in the brain unexplored. Astrocytes, the brain’s major glial cells, perform vital functions such as ion/energy homeostasis, neural plasticity regulation, and blood flow control.^24,25^ They play a crucial role to maintain brain homeostasis, and accumulating evidence also links astrocytes to aging and age-related neurodegenerative diseases, including their metabolic dysfunction and dysregulation of mitophagy.^26,27^ Despite existing measurements of neuronal mitophagy *ex vivo*, astrocytic mitophagy remains largely overlooked.

To address these knowledge gaps and technical challenges, we utilized two-photon imaging with mt-Keima probe in awake, behaving mice to uncover real-time mitophagy dynamics in layer 2/3 somatosensory cortex (SC). We quantified and compared mitophagy events in both neurons and astrocytes from early-aged and old-aged mice. We also used transmitted electron microscopy (TEM) to investigate the neuronal and astrocytic mitochondrial ultrastructure and identify mitophagy-like events. Finally, we assessed the effect of mitophagy induction through nicotinamide riboside (NR) administration in old-aged mice, revealing cell type-specific responses to therapeutic intervention.

## Results

### Calibration of two-photon laser excitation condition for *in vivo* mt-Keima imaging

To explore the real-time mitophagy in the mouse brain, the mt-Keima probe, which is a mitochondria-targeted mKeima,^20,28^ was used to illustrate mitophagy events in combination with two-photon imaging in awake behaving mice. The mKeima is a large Stokes shift fluorescent protein with a bimodal excitation spectrum: it peaks at a single-photon excitation wavelength of 440 nm in neutral pH environments (about pH 7) and shifts to approximately 590 nm under acidic conditions (about pH 4), with emission consistently peaking at 620 nm.^29,30^ This property allows for dual-excitation ratio imaging to detect autolysosome and mitolysosome formation and their related autophagy or mitophagy events, respectively.^20,28^ With two-photon excitation, previous studies have shown that under neutral pH condition the peak excitation wavelength of mKeima is at around wavelength of 880 - 900 nm, and the two-photon absorption (TPA) drops significantly below 800 nm and above 1000 nm, ^31,32^ however, no similar data was available under acid condition.

Since sequential two-photon scanning with two distinct excitation wavelengths is technically challenging for dynamic imaging in live, behaving mice, we sought the most suitable single excitation two-photon wavelength that maximizes mt-Keima signals from acidic environments (functional mitolysosomes) while minimizing mt-Keima signals from neutral environments. To achieve this, we used a HeLa cell line stably expressing mt-Keima.^20^ Given the limited laser power output and low TPA above 1000 nm, we tested wavelengths between 800 nm and 900 nm. Using 800 nm excitation wavelength, we achieved maximum mt-Keima fluorescent signals at pH 4 and minimal signals at pH 7 in these Hela cells (**Figure 1A)**. This wavelength effectively ensured that the majority of the captured signal originated from the acidic environment of active mitolysosomes. Thus, the 800 nm two-photon excitation wavelength was selected to perform following *in vivo* imaging experiments.

**Figure 1.**
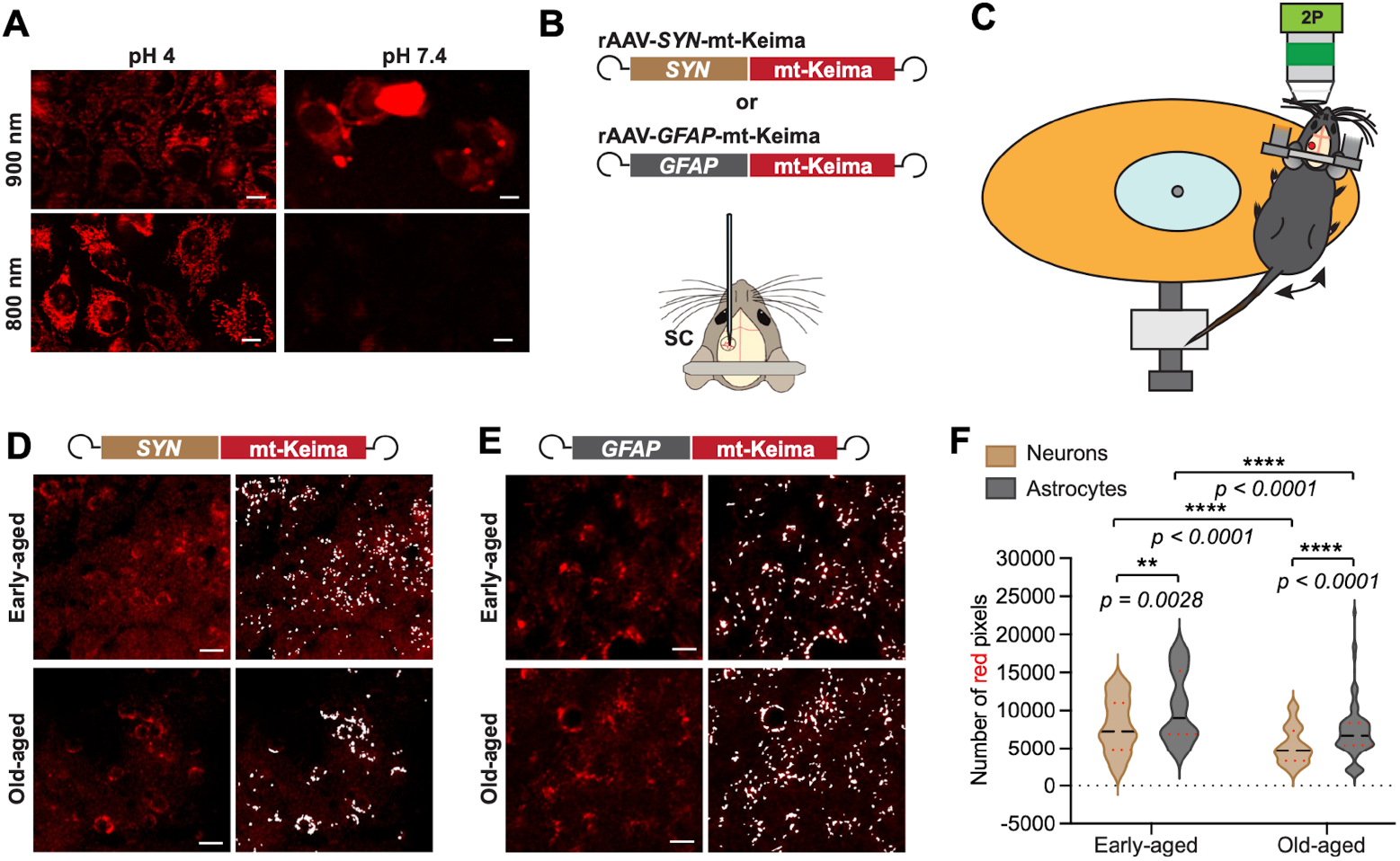
Two-photon mt-Keima imaging in neurons and astrocytes during aging. **(A)** Two-photon imaging of mt-Keima expressing Hela cell line under acidic (pH 4) and neutral (pH 7.4) environment with two-photon excitation wavelengths at 900 nm and 800 nm. Scale bar, 10 μm. **(B)** Illustration of rAAV constructs expressing mt-Keima under *SYN* and *GFAP* promoters, and schematic representation of rAAV delivery in mouse somatosensory cortex (SC). **(C)** Illustration of the two-photon (2P) imaging setup in head-fixed awake behaving mice. **(D-E)** Representative images (left side images in each group) of rAAV-*SYN*-mt-Keima **(D)** and rAAV-*GFAP*-mt-Keima **(E)** transduced SC in early-aged and old-aged wild-type mice. Signal extraction for quantitative analysis using *Ilastik* software were showing on the right in each group. Scale bar, 10 µm. **(F)** Quantitative comparison of detected mt-Keima signals indicating mitophagy levels in neurons and astrocytes during aging. Early-aged group, rAAV-*SYN*-mt-Keima, *n* = 60 from 3 mice; rAAV-*GFAP*-mt-Keima, *n* = 80 from 6 mice; old-aged group, rAAV-*SYN*-mt-Keima, *n* = 69 from 5 mice; rAAV-*GFAP*-mt-Keima, *n* = 94 from 3 mice. Statistical significance was tested by the Mann-Whitney test. Medians and quartiles are indicated by black and red dotted lines, respectively; ns, non-significant; *p* < 0.05 (*); *p* < 0.01 (**); *p* < 0.001 (***); *p* < 0.0001 (****).

### Neurons and astrocytes exhibited distinct mitophagy dynamics during aging

To measure mitophagy levels in both neurons and astrocytes, *SYN* and *GFAP* promoters were used to drive expression of mt-Keima in neurons and astrocytes via recombinant adeno-associated virus (rAAV) gene delivery, respectively (**Figure 1B**), and both promoters show cell type-specificity of neurons and astrocytes in mouse SC region (**Figure S1A**). To reveal mitophagy levels in neurons and astrocytes during aging, two weeks after rAAV transduction, both early-aged (2-3-month-old) and old-aged (18-20-month-old) adult wild-type mice expressing mt-Keima in either neurons or astrocytes in layer 2/3 SC were placed on a freely spinning disc for imaging (**Figure 1C-E, Video S1 - S4**). We quantified mitophagy levels based on the mt-Keima signals acquired from layer 2/3 SC neurons or astrocytes (**Figure 1D-E**, right panels). Our data revealed two key findings regarding mitophagy dynamics (**Figure 1F**). Firstly, we observed that mitophagy levels decreased significantly with age in both cell types. Old-aged mice exhibited approximately a 34% reduction in neuronal mitophagy and a 31% reduction in astrocytic mitophagy compared to the early-aged group. Secondly, in both age groups, astrocytes consistently maintained significantly higher mitophagy levels than neurons (approximately 31% higher in early-aged and 36% higher in old-aged mice). Collectively, our *in vivo* characterization of neuronal and astrocytic mitophagy in the live mouse SC suggests that these cell types follow two distinct dynamic patterns during the aging process.

### Aging affected mitochondrial features in neurons and astrocytes

To complement our two-photon imaging data, we used traditional TEM to assess mitochondrial ultrastructure in the SC regions of the same early- and old-aged mice (**Figure 2A and 2B**). We quantified mitophagy/autophagy-like events, total mitochondrial number, percentages of damaged mitochondria (characterized by loss of cristae and/or double membranes), and morphological details (size, diameter, perimeter) in both neurons and astrocytes. To ensure cell-type specificity in TEM, astrocytic quantification was limited strictly to endfeet in direct contact with blood vessels (**Figure 2B**). We did not observe significant differences in mitophagy/autophagy-like events using TEM in both neurons and astrocytic endfeet (**Figure 2C and 2D**). However, quantification on various mitochondrial features revealed age- and cell type-related changes within regions of interest (ROIs). During aging, the total number of mitochondria and the percentage of damaged mitochondria increased in neurons (**Figure 2E and 2F**), and neuronal mitochondrial areas and diameters also became significantly larger (**Figure 2G**), suggesting age-dependent morphological changes in neurons. In astrocytic endfeet, we observed only an increase in the percentage of damaged mitochondria during aging, while the total number of mitochondria and morphological details remained similar to early-aged mice (**Figure 2E and 2F**). Regardless of age groups, the astrocytic mitochondrial areas, diameters and perimeters were all significantly larger than those in neurons (**Figure 2G, Table 1**), which reflect an age-independent, cell-type specific mitochondrial morphological characteristics in mouse SC.

**Table 1.** Mitochondrial parameter comparison in neurons and astrocytes with TEM.

|  | **Mitochondrial parameter** | **Neurons**<br>**(mean ± SEM)** | **Astrocytes**<br>**(mean ± SEM)** | ***p*-value**<br>**(Mann-Whitney test)** |
| --- | --- | --- | --- | --- |
| **Early-aged group** | Area | 0.068 ± 0.001 | 0.128 ± 0.006 | *p* < 0.001 |
|  | Diameter | 0.209 ± 0.001 | 0.245 ± 0.004 | *p* < 0.001 |
|  | Perimeter | 0.986 ± 0.009 | 1.506 ± 0.052 | *p* < 0.001 |
| **Old-aged group** | Area | 0.069 ± 0.001 | 0.122 ± 0.007 | *p* < 0.001 |
|  | Diameter | 0.214 ± 0.001 | 0.261 ± 0.007 | *p* < 0.001 |
|  | Perimeter | 0.994 ± 0.011 | 1.380 ± 0.053 | *p* < 0.001 |

**Figure 2.**
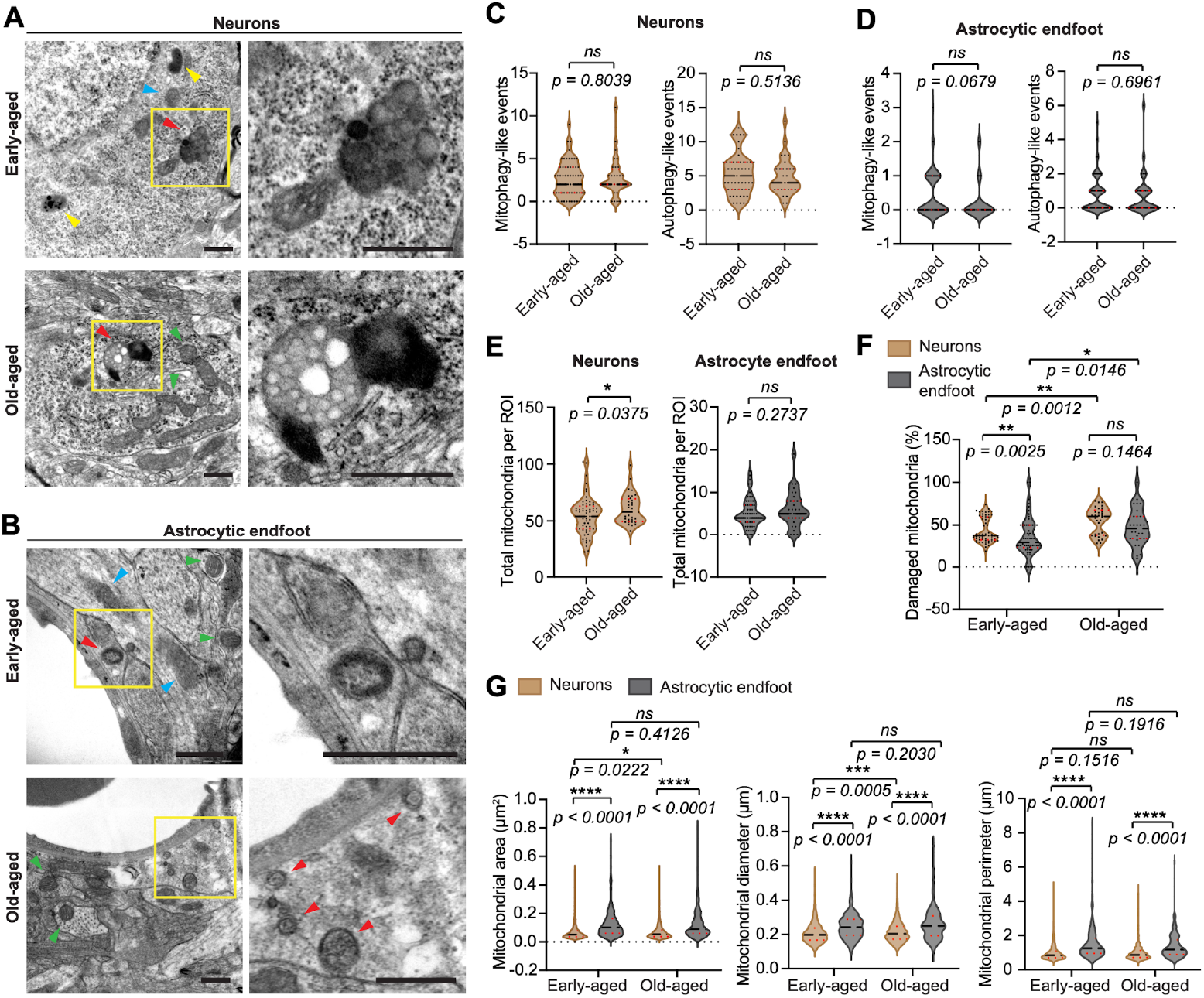
TEM analyses on mitochondrial features during aging. Representative TEM images of neurons **(A)** and astrocytic endfeet abutting the blood vessels **(B)** in early-aged and old-aged wild-type mice. Green arrowheads indicate healthy mitochondria, blue arrowheads indicate damaged mitochondria, yellow arrowheads show structures of lipofuscin, red arrowheads show mitophagy-like events. Scale bars, 500 nm. **(C)** Comparison of total numbers of mitochondria in neurons (Welch’s t-test) and astrocytic endfeet during aging. **(D)** Comparison of percentages of damaged mitochondria during aging in neurons and astrocytic endfeet. Neurons vs astrocytic endfeet in old-aged group, Welch’s t-test. Neurons, early-aged, *n* = 63 from 6 mice; old-aged, *n* = 32 from 3 mice; astrocytic endfeet, early-aged, *n* = 57 from 6 mice, old-aged, *n* = 28 from 3 mice. **(E-F)** Comparison of mitophagy-like and autophagy-like events in neurons **(E)** and astrocytic endfeet **(F)** between two age groups. Neurons, early-aged, *n* = 63 from 6 mice, old-aged, *n* = 32 from 3 mice; astrocytes, early-aged, *n* = 60 from 6 mice, old-aged, *n* = 31 from 3 mice. **(G)** Comparison of mitochondrial areas, diameter and perimeter during aging in neurons and astrocytes. Neurons, early-aged, *n* = 2971 from 6 mice; old-aged, *n* = 1710 from 3 mice; astrocytic endfeet, early-aged, *n* = 307 from 6 mice; old-aged, *n* = 182 from 3 mice. Statistical significance was tested by Mann-Whitney test, unless stated otherwise. Medians and quartiles are indicated by black and red dotted lines, respectively; ns, non-significant; *p* < 0.05 (*); *p* < 0.01 (**); *p* < 0.001 (***); *p* < 0.0001 (****).

### NR treatment improved neuronal and astrocytic mitophagy in old-aged mice

To further apply the two-photon mt-Keima *in vivo* imaging to a therapeutic context, we investigated a pharmacological activation of mitophagy using NR. NR is known to enhance mitophagy by increasing nicotinamide adenine dinucleotide (NAD^+^) levels, which in turn activates NAD^+^-dependent pathways involved in mitochondrial quality control.^33^ We aimed to determine whether NR could ameliorate the age-dependent decline of brain mitophagy *in vivo*. Old-aged mice previous imaged were treated with 12 mM NR for 3 weeks and subsequently re-imaged using two-photon microscopy (**Figure 3A**). The NR treatment significantly increased mitophagy levels in both neurons (56% improvement) and astrocytes (19% improvement), indicating an enhancement in mitochondrial removal mechanisms (**Figure 3B**). However, subsequent TEM analysis of the SC tissues showed that the NR treatment did not significantly alter mitophagy-like/autophagy-like events (**Figure 3C and 3D**), the total number of mitochondria or the percentage of damaged mitochondria in both neurons and astrocytic endfeet when compared to untreated conditions (**Figure 3E and 3F**). However, neuronal mitochondrial sizes measured by area and perimeter were significantly increased after the NR treatment, whereas those in astrocytic endfeet did not (**Figure 3G**), suggesting changes in neuronal mitochondrial morphological characteristics following NR intervention.

**Figure 3.**
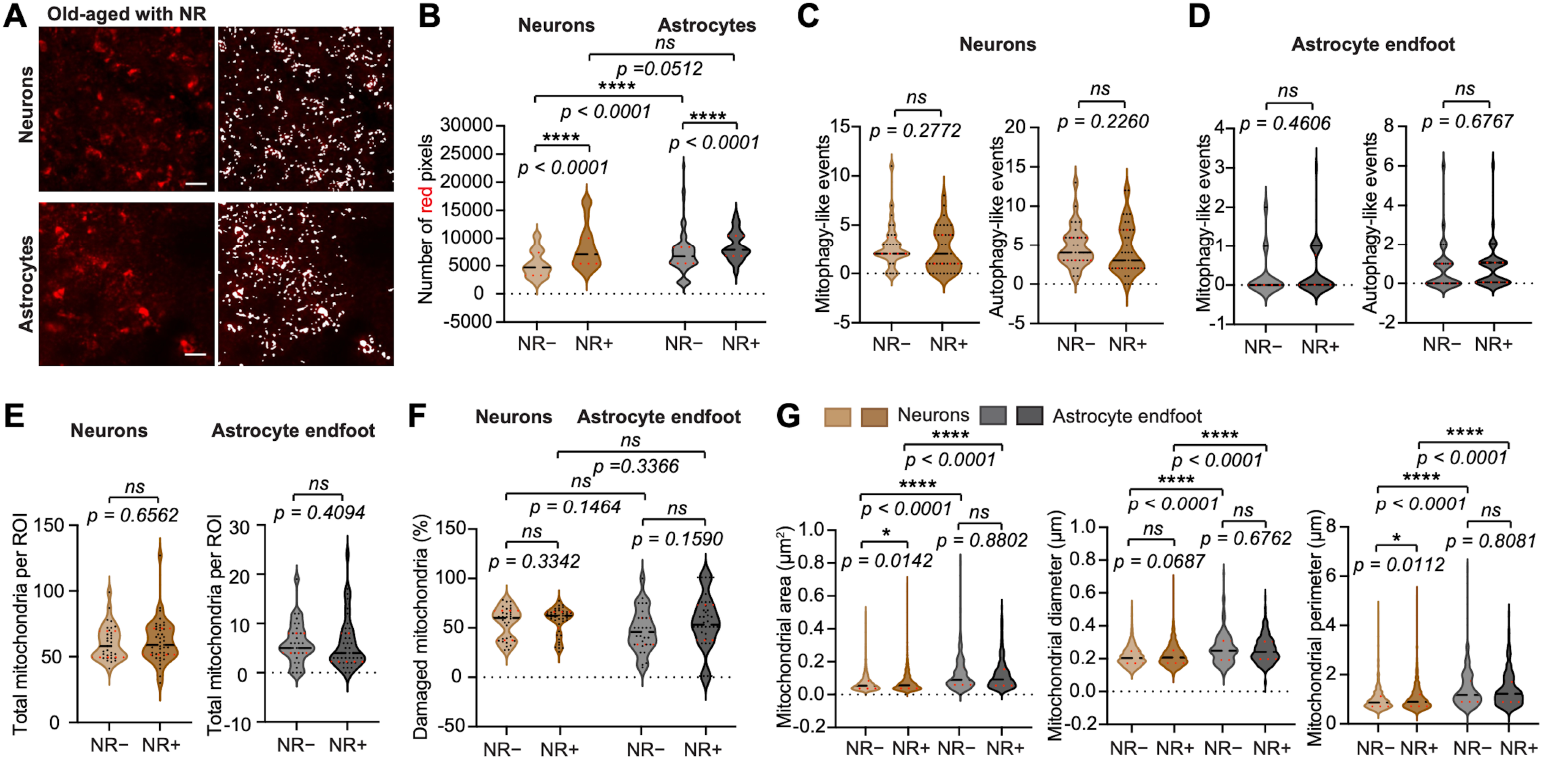
Nicotinamide riboside (NR) treatment effects on neurons and astrocytes in old-aged mice. **(A)** Representative images of rAAV-*SYN*-mt-Keima and rAAV-*GFAP*-mt-Keima transduced neurons and astrocytes, respectively, in SC in old-aged adult mice with NR treatment (left images). Signal extraction for quantitative analysis using *Ilastik* software were showing on the right. Scale bar, 10 µm. **(B)** Quantitative comparison of detected mt-Keima signals indicating mitophagy levels in neurons and astrocytes before and after NR treatment. Group without NR, rAAV-*SYN*-mt-Keima, *n* = 69 from 5 mice; rAAV-*GFAP*-mt-Keima, *n* = 94 from 3 mice; group with NR, rAAV-*SYN*-mt-Keima, *n* = 65 from 5 mice; rAAV-*GFAP*-mt-Keima, *n* = 90 from 3 mice. **(C-D)** Comparison of mitophagy-like events and autophagy-like in neurons **(C)** and astrocytic endfeet **(D)** before and after NR treatment. Neurons, group without NR, *n* = 32 from 3 mice, group with NR, *n* = 41 from 4 mice; astrocytic endfeet, group without NR, *n* = 31 from 3 mice, group with NR, *n* = 40 from 4 mice. **(E)** Comparison of total numbers of mitochondria in neurons and astrocytic endfeet before and after NR treatment. **(F)** Comparison of percentages of damaged mitochondria in neurons and astrocytic endfeet before and after NR treatment. Neurons vs astrocytic endfeet in old-aged group, Welch’s t-test. Neurons, group without NR, *n* = 32 from 3 mice, group with NR, *n* = 41 from 4 mice; astrocytic endfeet, group without NR, *n* = 28 from 3 mice, group with NR, *n* = 38 from 4 mice. **(G)** Comparison of mitochondrial areas, diameter and perimeter in neurons and astrocytes before and after NR treatment. Neurons, group without NR, *n* = 1710 from 3 mice; group with NR, *n* = 2295 from 4 mice; astrocytic endfeet, group without NR, *n* = 182 from 3 mice; group with NR, *n* = 231 from 4 mice. Statistical significance was tested by Mann-Whitney test, unless stated otherwise. Medians and quartiles are indicated by black and red dotted lines, respectively; ns, non-significant; *p* < 0.05 (*); *p* < 0.01 (**); *p* < 0.001 (***); *p* < 0.0001 (****).

## Discussion

Aging is characterized by a progressive decline in cellular function, with both mitochondrial dysfunction and impaired autophagy emerging as central hallmarks in research concerning brain aging and age-related neurodegenerative diseases.^1,34,35^ In this study, we present the first real-time *in vivo* two-photon imaging assessment of cell type-specific mitophagy in the SC of early-aged and old-aged adult mice. Our findings revealed distinct patterns of mitophagy dynamics in neurons and astrocytes during the aging process. Specifically, we observed that mitophagy levels, as measured by mt-Keima reporter in the SC, were significantly reduced with age in both cell types. Crucially, astrocytic mitophagy levels were consistently higher than neuronal levels in both age groups (**Figure 1F**). These data underscore the need for more comprehensive approaches to study brain metabolism during aging that account for distinct contributions of both neuronal and glial populations.

This study offered a significant technical advantage, providing higher accuracy in describing the real-time dynamics of mitophagy in the brain compared to previously published research. The employment of two-photon microscopy has fundamentally improved *in vivo* experimental conditions of using mitophagy reporters. When compared to imaging of mt-Keima in acute brain slices^20^ or confocal imaging of mt-QC in the fixed transgenic mouse brain,^22,36^ our approach restored the real-time mitophagy dynamics in living mouse brains. To our surprise, we did not observe fast temporal dynamics of mitophagy during imaging. Instead, the recorded mitophagy pattern remained relatively stable in both neurons and astrocytes over five-minute recording period (**Video S1 - S4**).

In this study, the rAAV delivery of the mitophagy reporter, the mt-Keima probe, has achieved cell type-specific expression that has not been very clearly illustrated in previous studies using transgenic mice. Transgenic mt-QC mice typically use a CAG promoter, which primarily drives expression in neurons,^19,22,36^ but also may result in expression in astrocytes and oligodendrocytes.^37^ While Schmid et al. demonstrated neuronal specific expression of mt-QC in *Drosophila*,^38^ their analysis was performed on fixed brain tissue. Direct comparison between the results from two species and *in vivo* versus *ex vivo* experimental conditions is challenging. Thus, our results on measuring mitophagy levels using two-photon microscopy in the SC provide a valuable perspective on cell type specificity of brain mitophagy.

While the use of mt-Keima allows for real-time *in vivo* mitophagy assessment, it does not provide information on status of healthy and impaired mitochondria. Our TEM analyses, offering subcellular resolution, helped to clarify this property. The TEM results showed that both the total count of neuronal mitochondria and the percentage of damaged neuronal and astrocytic endfoot mitochondria were all increased during aging (**Figure 2E and 2F**, respectively). As previously reported in the *Drosophila* brain,^38^ this also suggests that mitochondrial quality control is a more critical determinant of brain aging than the simple loss of mitochondrial content. Meanwhile, we also observed that the actual size of mitochondria in astrocytic endfeet was larger than in neurons (**Figure 2G**). Functionally, astrocytes possess several microcompartments with distinct activation responses, as measured by changes in intracellular calcium concentration.^39,40^ Astrocytic endfeet are one distinct functional microcompartments that participates in forming the blood-brain barrier, and their specific metabolic demands might differ from mitochondria in other microcompartments of astrocytes. Although we should not generalize our findings in the defined endfoot compartment to the entire astrocyte, these data, combined with the *in vivo* two-photon mt-Keima imaging, suggest that the distribution of mitochondria and metabolic demands likely differ between neurons and astrocytes, opening new avenues to understand the contribution of spatial and temporal changes in mitochondria during aging.

Furthermore, we have demonstrated that pharmacological activation of brain mitophagy with NR elevates mitophagy levels in both neurons and astrocytes in layer 2/3 in the mouse SC. The beneficial effects of NAD^+^ repletion in neurons have previously been shown in aging and disease models, resulting in notable increases in mitophagy and mitochondrial function.^41–43^ While it is hard to predict the precise mechanistic basis for these observed effects of NR treatment,, potential factors may involve NAD^+^ metabolism, lysosomal function, or mitophagy regulatory pathways in neurons and astrocytes.^44,45^ Follow-up mechanistic studies are urgently needed to fully understand the treatment effect of elevating mitophagy levels via this intervention.

## Limitations of the study

Due to the technical limitations of two-photon microscopy, traditional ratiometric imaging requiring two distinct excitation wavelengths was not feasible. Instead, we used an optimal single two-photon excitation wavelength of 800 nm, which maximized mt-Keima signals in acid environment while minimizing signals at neutral pH (**Figure 1A**). Given that the mt-Keima signals detected at pH 7.4 were very low, and recordings were performed under similar conditions, we considered 800 nm excitation suitable for measuring *in vivo* mt-Keima signals under acid conditions. Interestingly, we observed a previous unreported change in the mt-Keima emission spectrum when exciting with 800 nm versus 900 nm wavelengths. With higher power output at 800 nm compared to 900 nm from the two-photon laser, signals collected in the green emission channel at pH 4 and pH 7.4 were stronger at 900 nm excitation (**Figure S1B**). This indicates an emission spectrum shift when changing excitation wavelength, which has not been reported from previous studies with mt-Keima using both single photon and two-photon excitation.^29–32^ A further technical limitation is that when imaging the miniature structure of mitochondria, the spatial resolution obtained from two-photon microscopy was relatively poor when compared to previous studies using stimulated emission depletion microscopy,^20^ and individual mitochondria were not able to be illustrated. Additionally, the limited imaging depth of our method restricted our access to neurons and astrocytes in layer 2/3 of the SC. Therefore, the applicability of our conclusions to other brain regions, particularly the subcortical areas, required further investigation.

In addition, our TEM analyses did not fully recapitulate the two-photon imaging results. This may be due to the small sample size used for TEM (3-6 mice per group, approximately 10 images per cell type per mouse), which limited the statistical power to detect subtle differences. This limitation was particularly evident for the analysis intended for astrocytes, as we were restricted to performing TEM analysis solely on astrocytic endfeet. Compared to TEM study, the two-photon recording covered much larger brain areas. Expanding the TEM study cohort and incorporating functional assays of mitochondrial activity would strengthen these conclusions.

Several key questions remain unanswered and require future investigation. First, our data reported were all from male mice, and it is crucial to assess whether gender differences exist in mitophagy regulation and in response to NR treatment. Notably, previous research has reported sex-dependent mitophagy differences in cerebellar astrocytes, with higher mitophagy in females during midlife;^22^ similarly, there were sex- and region-specific shifts in chaperone-mediated autophagy with aging.^46^ Second, while we explored changes in mitophagy in neurons and astrocytes during aging in live mice, we did not include microglia. Third, building on the success of this study, *in vivo* two-photon microscopy-based mitophagy studies should be extended to other brain regions of live mice. Lastly, while our study focused primarily on structural and morphological changes, the functional implications of the astrocytic mitophagy decline remain unclear. Future study should examine whether age-related astrocytic mitophagy deficits directly contribute to any neuronal dysfunction, synaptic loss, or even cognitive decline. Similarly, cell type-specific functional studies on mitochondria treated with NR are still lacking and will be crucial for understanding its potential as a therapeutic agent.

## Acknowledgement

We thank Prof. Nuo Sun and Prof. Toren Finkel for providing the mt-Keima-based cell line with consent from the original owner.

## Author contributions

CRediT: **Beatriz Escobar-Doncel**: Formal analysis, Methodology, Visualization, Writing – original draft, Writing – review & editing; **Xiaoyi Zhang**: Investigation, Methodology; **Mario Fernández de la Puebla**: Data curation, Methodology; **Shreyas Balachandra Rao**: Investigation, Methodology; **Synnøve Algrøy Fjeldstad**: Investigation; **Maja Tvedt Dahle**: Investigation; **Mahmood Amiry-Moghaddam**: Methodology; **Magnar Bjørås**: Methodology, Funding acquisition; **Evandro Fei Fang**: Conceptualization, Funding acquisition, Supervision, Writing – review & editing; **Wannan Tang**: Conceptualization, Funding acquisition, Resources, Project administration, Supervision, Visualization, Writing – original draft, Writing – review & editing.

## Data availability statement

### Lead contact

Information and requests for resources and reagents all be directed to and will be fulfilled by the lead contact Wannan Tang.

### Materials availability

This study did not generate new, unique reagents.

### Data and code availability

The original recorded data are available upon requests by the lead contact Wannan Tang. All original code for image pre-processing has been deposited at GitHub and is publicly available at https://doi.org/10.1016/j.jneumeth.2017.07.031. The mitophagy quantification was performed using an open-source image classification and segmentation software Ilastik (www.ilastik.org).

## STAR Methods

### Animals

Male mice with age of 2-3-month-old (early-aged) and 18-20-month-old (old-aged) were housed with a 12-hour light/dark cycle (light on at 8 a.m.) with running wheels and cardboard houses et to enrich the housing environment. All animal handling and experimental procedures were performed according to the European Animal Research law (Directive 2010/63/EU) and approved by the Norwegian Food Safety Authority (project number: FOTS #27090 and #29730).

### Nicotinamide Riboside (NR) treatment

After two-photon imaging, all old-aged mice were administrated with NR (12 mM NR in drinking water) for a duration of 3 weeks. To ensure consistency and maintain NR stability, drinking water was replaced once per week in both groups throughout the treatment period.

### Plasmid construction and virus production

The DNA sequence of the mt-Keima^20^ was synthesized and subcloned in to the recombinant adeno-associated virus (rAAV) vector pAAV-6P-SEWB^47^ and pAAV-GFAP-GCaMP5E^39^ with 5’ BamHI and 3’ HindIII to create pAAV-*SYN*-mt-Keima and pAAV-*GFAP*-mt-Keima, respectively. Serotype 2/1 rAAVs from pAAV-*SYN*-mt-Keima and pAAV-*GFAP*-mt-Keima were produced and purified by AVB Sepharose affinity chromatography,^48^ following titration with real-time PCR (rAAV titer about 1.0 – 6.0 x 10^12^ viral genomes/mL, TaqMan Assay, Applied Biosystems).

### Surgical procedures and virus transduction

Mice were briefly anesthetized with 3% isoflurane in an induction chamber and then quickly fastened on a stereotaxic frame with a nose cone (Model 963, KOPF Instruments). Animals were maintained under anesthesia with 1.2-1.5% isoflurane in a mixture of 80% oxygen and 20% room air and buprenorphine 0.1 mg/kg s.c. preemptively for analgesia. Before the surgery, Bupivacaine was administered subcutaneously over the skull and left for 10 min. Skin and connective tissues over the somatosensory cortex were removed. A custom-made titanium head-bar was glued to the mouse skull using cyanoacrylate glue (Loctite-super glue) and sealed with dental cement (Meliodent, Kulter). A 2.5-mm-diameter craniotomy was then drilled with its center coordinates relative to Bregma of anteroposterior (AP) −1.1 mm and mediolateral (ML) 3.5 mm. A fine dental drill was used to carefully cut the skull with intermittent soaking and removal of debris until only ~0.1 mm of the bone thickness was left. After 10 min of soaking in 0.9% saline, the bone flap was carefully removed. The rAAVs were slowly injected (20-30 nL/minute) at three evenly spaced locations (70 nL each at 400 μm depth) positioned to stay clear from large blood vessels. A glass plug consisting of two coverslips with diameters of 2.5 and 3.5 mm glued together with ultraviolet curing glue (Norland Optical Adhesive No. 61) was placed and fixed on the craniotomy using cyanoacrylate-gel glue (Loctite-super glue gel) and reinforced with dental cement (Meliodent, Kulter). The glass window was slightly pressing the dura to prevent dural overgrowth. Mice with implant complications were excluded from the further experiments.

### Two-photon imaging

Three to seven weeks after the surgery, mice were imaged under a two-photon microscope (Ultima IV, Bruker/Prairie Technologies) with a 16 × 0.8 NA water-immersion objective (model CFI75 LWD 16XW, Nikon), an InSight DeepSee laser (Spectra-Physics) and Peltier-cooled photomultiplier tubes (model 7422PA-40, Hamamatsu Photonics K.K). Optical filter settings: a main dichroic filter (ZT473-488/594/NIRtpc) in the detection unit reflects the collected light. The light enters the detector house with four channels, passing a ZET473-488/594/NIRm filter to shield the photo multiplier tubes from the reflective light. Inside the detector house, the light is split into two fractions separated at the wavelength of 560 nm by a dichroic filter (T560lpxr). The green wavelength light is further guided by a secondary dichroic beam splitter at 495 nm (T495lpxr) and filtered by a ET525/50m-2p bandpass filter. And the red wavelength light is similarly guided by a secondary beam splitter at 640 nm (T640lpxr) and subsequently filtered by a ET595/50m-2p bandpass filter. The emitted photons were detected with Peltier cooled photomultiplier tubes (model 7422PA-40 by Hamamatsu Photonics K.K.). After calibration, the excitation wavelength of 800 nm was used to record mt-Keima signals. Images with 512×512 pixels resolution were acquired at about 1 Hz. Imaging time series of 5 minutes were recorded from multiple field of views (FOVs) in in layer 2/3 of somatosensory cortex of each animal.

### Imaging analysis

Imaging raw data were corrected for motion artifacts using the NoRMCorre movement correction algorithm ^49^. Recordings that could not be sufficiently motion-corrected were excluded from the analysis. Subsequently, the stabilized T-series were imported into ImageJ for further processing. From each T-series, sets of five consecutive frames were stacked to generate one average Z-projection image, corresponding to 5 seconds of recording time (1 Hz), with inclusion criteria requiring uninterrupted signal quality (i.e. noise recording) across all five frames. Up to five sets (5-frame stack) were selected per stabilized T-series with one minute time internals from the recording, ensuring a complete representation of a full recording. The same sets were selected for both channels to guarantee consistency. Each five-frame stack was then converted into a single image by calculating the average intensity projection. To minimize edge artifacts (e.g. noise recording) and ensure consistent image dimensions across the dataset, all images were uniformly cropped, irrespective of the presence of visible artifacts.

### mt-Keima analysis by Ilastik

All images were further processed using Ilastik, an open-source image classification and segmentation software (www.ilastik.org).^50^ Ilastik applies supervised machine learning algorithms to rapidly quantify image features, enabling the establishment of consistent thresholding parameters across all images. Feature selection included color/intensity (σ = 0.3, 0.7, 1.00, 1.60, 3.50, 5.00, and 10.00), edge (σ = 0.7, 1.6, and 5.00), and texture (σ = 1.00, 3.50, and 10.00). Pixel classification was then performed to distinguish background noise (‘background’ label) from true signal (‘pixel’ label). Up to five sets of images per condition were used to train the classifier. Thresholding for the ‘pixel’ label was applied using the simple method, with smoothing parameters set to 0.0 for both minimum and maximum values, and a threshold value of 0.45. A size filter was also applied, with the minimum object size set to 9 pixels, corresponding approximately to the size of a mitochondrion. Following training and thresholding, experimental images were batch-processed, and results were manually inspected. After the batch processing, object information was exported for downstream quantification, including total pixel number, object count, and object feature analysis. These outputs were then used for subsequent ratiometric quantification.

### Sample preparation and embedding for TEM

For perfusion and tissue preparation, mice from all groups were deeply anesthetized using 5% isoflurane and then were transcardially perfusion fixed, initially using 2% dextran in 0.1 M phosphate buffer for 20 sec followed by the fixative for 20 min. The fixative consisted of 4% formaldehyde and 0.1% glutaraldehyde in 0.1 M phosphate buffer at pH 7.4. Following perfusion, the brains were carefully dissected out, post fixed overnight in the fixative. The following day, the brains were stored in 1:10 dilution of the fixative in 0.1 M phosphate buffer until further use. The samples were embedded for electron microscopy as follows: small blocks of volume ~1 mm^3^ from tissue were dissected out from the fixed brains and were subjected to freeze substitution procedure as described previously ^51^. Briefly, tissue blocks were cryoprotected in 4% glucose for overnight, followed by suspension in graded glycerol solution (10%, 20% and 30% glycerol in 0.1 M phosphate buffer) for 30 min in each gradient, with the final step being overnight suspension in 30% glycerol. The following day, tissues were rapidly frozen in propane cooled to −170 °C using liquid nitrogen before subjecting to freeze substitution. Samples were later embedded in methacrylate resin (Lowicryl HM20) and polymerized by UV irradiation at −45 °C for 48 hours. Ultrathin sections of thickness 80-90 nm were cut and placed on formvar carbon coated support film in Ni-grids and counterstained using 2% uranyl acetate and 0.3% lead citrate for 90 sec each. Images were acquired using a Tecnai 12 electron microscope (FEI) at 80 kV.

### TEM analysis and quantification

TEM images of neurons and astrocytic endfeet were analyzed for the quantification of mitochondria number and classification, mitophagy and autophagy events. The region of interest (ROI) for neurons included soma, synapses, dendrites and axons, and for astrocytes included endfeet that abut the blood vessels. The general approach for TEM analysis of the brain was based on previously published references ([Introduction » Fine Structure of the Aging Brain | Boston University](https://www.bu.edu/agingbrain/introduction/)). Mitochondria were classified based on structural criteria: mitochondria with a double membrane and clearly visible cristae were considered *healthy*; those displaying a double membrane without evident cristae were categorized as *damaged*; and structures where these features could not be confidently identified were labelled as *unrecognized*. Quantification of mitophagy and autophagy events were performed according to established protocols.^52–54^ The mitophagy rate was calculated as the total number of mitophagy events per ROI divided by the total number of mitochondria per ROI.

### Statistics

Statistical analysis was performed using Prism (Version 9.3.1 for Mac OSX, GraphPad Software). Details about the choice of statistics are described in the main text and figure legends. Full descriptions of statistical parameters were accessed with the original data before choosing a suitable analysis.

## Supplemental information

Figure S1, related to Figure 1.

Video S1 – S4, related to Figure 1.

## Abbreviations

NAD^+^: nicotinamide adenine dinucleotide
NR: nicotinamide riboside
ROI: region of interest
SC: somatosensory cortex
TEM: transmitted electron microscopy
TPA: two-photon absorption

## References

1. Mattson, M.P., and Arumugam, T. V. (2018). Hallmarks of Brain Aging: Adaptive and Pathological Modification by Metabolic States. Cell Metab. 27, 1176–1199. 10.1016/j.cmet.2018.05.011.

2. Caponio, D., Veverová, K., Zhang, S., Shi, L., Wong, G., Vyhnalek, M., and Fang, E.F. (2022). Compromised autophagy and mitophagy in brain ageing and Alzheimer’s diseases. Aging Brain 2, 100056. 10.1016/j.nbas.2022.100056.

3. Bartman, S., Coppotelli, G., and Ross, J.M. (2024). Mitochondrial Dysfunction: A Key Player in Brain Aging and Diseases. Curr. Issues Mol. Biol. 46, 1987–2026. 10.3390/cimb46030130.

4. Onishi, M., Yamano, K., Sato, M., Matsuda, N., and Okamoto, K. (2021). Molecular mechanisms and physiological functions of mitophagy. EMBO J. 40, 1–27. 10.15252/embj.2020104705.

5. Fang, E.F., Hou, Y., Palikaras, K., Adriaanse, B.A., Kerr, J.S., Yang, B., Lautrup, S., Hasan-Olive, M.M., Caponio, D., Dan, X., et al. (2019). Mitophagy inhibits amyloid-β and tau pathology and reverses cognitive deficits in models of Alzheimer’s disease. Nat. Neurosci. 22, 401–412. 10.1038/s41593-018-0332-9.

6. Lautrup, S., Sinclair, D.A., Mattson, M.P., and Fang, E.F. (2019). NAD+ in Brain Aging and Neurodegenerative Disorders. Cell Metab. 30, 630–655. 10.1016/j.cmet.2019.09.001.

7. Sun, N., Youle, R.J., and Finkel, T. (2016). The Mitochondrial Basis of Aging. Mol. Cell 61, 654–666. 10.1016/j.molcel.2016.01.028.

8. Kerr, J.S., Adriaanse, B.A., Greig, N.H., Mattson, M.P., Cader, M.Z., Bohr, V.A., and Fang, E.F. (2017). Mitophagy and Alzheimer’s Disease: Cellular and Molecular Mechanisms. Trends Neurosci. 40, 151–166. 10.1016/j.tins.2017.01.002.

9. Kobro-Flatmoen, A., Lagartos-Donate, M.J., Aman, Y., Edison, P., Witter, M.P., and Fang, E.F. (2021). Re-emphasizing early Alzheimer’s disease pathology starting in select entorhinal neurons, with a special focus on mitophagy. Ageing Res. Rev. 67, 101307. 10.1016/j.arr.2021.101307.

10. Rodger, C.E., McWilliams, T.G., and Ganley, I.G. (2018). Mammalian mitophagy – from in vitro molecules to in vivo models. FEBS J. 285, 1185–1202. 10.1111/febs.14336.

11. Ashrafi, G., and Schwarz, T.L. (2013). The pathways of mitophagy for quality control and clearance of mitochondria. Cell Death Differ. 20, 31–42. 10.1038/cdd.2012.81.

12. Picca, A., Faitg, J., Auwerx, J., Ferrucci, L., and D’Amico, D. (2023). Mitophagy in human health, ageing and disease. Nat. Metab. 5, 2047–2061. 10.1038/s42255-023-00930-8.

13. Li, Z., Wu, Q., Liu, L., Sun, S., Sun, S., Wang, Z., and Li, J. (2021). Determination of mitophagy by electron microscope. Methods Cell Biol. 165, 103–110. 10.1016/bs.mcb.2020.10.015.

14. Hernandez, G., Thornton, C., Stotland, A., Lui, D., Sin, J., Ramil, J., Magee, N., Andres, A., Quarato, G., Carreira, R.S., et al. (2013). MitoTimer: A novel tool for monitoring mitochondrial turnover. Autophagy 9, 1852–1861. 10.4161/auto.26501.

15. Dolman, N.J., Chambers, K.M., Mandavilli, B., Batchelor, R.H., and Janes, M.S. (2013). Tools and techniques to measure mitophagy using fluorescence microscopy. Autophagy 9, 1653–1662. 10.4161/auto.24001.

16. Goldsmith, J., Ordureau, A., Harper, J.W., and Holzbaur, E.L.F. (2022). Brain-derived autophagosome profiling reveals the engulfment of nucleoid-enriched mitochondrial fragments by basal autophagy in neurons. Neuron 110, 967-976.e8. 10.1016/j.neuron.2021.12.029.

17. Kallergi, E., Siva Sankar, D., Matera, A., Kolaxi, A., Paolicelli, R.C., Dengjel, J., and Nikoletopoulou, V. (2023). Profiling of purified autophagic vesicle degradome in the maturing and aging brain. Neuron 111, 2329-2347.e7. 10.1016/j.neuron.2023.05.011.

18. Sargsyan, A., Cai, J., Fandino, L.B., Labasky, M.E., Forostyan, T., Colosimo, L.K., Thompson, S.J., and Graham, T.E. (2015). Rapid parallel measurements of macroautophagy and mitophagy in mammalian cells using a single fluorescent biosensor. Sci. Rep. 5, 1–11. 10.1038/srep12397.

19. McWilliams, T.G., Prescott, A.R., Allen, G.F.G., Tamjar, J., Munson, M.J., Thomson, C., Muqit, M.M.K., and Ganley, I.G. (2016). Mito-QC illuminates mitophagy and mitochondrial architecture in vivo. J. Cell Biol. 214, 333–345. 10.1083/jcb.201603039.

20. Sun, N., Yun, J., Liu, J., Malide, D., Liu, C., Rovira, I.I., Holmström, K.M., Fergusson, M.M., Yoo, Y.H., Combs, C.A., et al. (2015). Measuring In Vivo Mitophagy. Mol. Cell 60, 685–696. 10.1016/j.molcel.2015.10.009.

21. Sun, N., Malide, D., Liu, J., Rovira, I.I., Combs, C.A., and Finkel, T. (2017). A fluorescence-based imaging method to measure in vitro and in vivo mitophagy using mt-Keima. Nat. Protoc. 12, 1576–1587. 10.1038/nprot.2017.060.

22. Rappe, A., Vihinen, H.A., Suomi, F., Hassinen, A.J., Ehsan, H., Jokitalo, E.S., and McWilliams, T.G. (2024). Longitudinal autophagy profiling of the mammalian brain reveals sustained mitophagy throughout healthy aging. EMBO J. 43, 6199–6231. 10.1038/s44318-024-00241-y.

23. Zimmermann, A., Madeo, F., Diwan, A., Sadoshima, J., Sedej, S., Kroemer, G., and Abdellatif, M. (2024). Metabolic control of mitophagy. Eur. J. Clin. Invest. 54, 1–19. 10.1111/eci.14138.

24. Bazargani, N., and Attwell, D. (2016). Astrocyte calcium signaling: The third wave, 10.1038/nn.4201 https://doi.org/10.1038/nn.4201.

25. Santello, M., Toni, N., and Volterra, A. (2019). Astrocyte function from information processing to cognition and cognitive impairment, 10.1038/s41593-018-0325-8 https://doi.org/10.1038/s41593-018-0325-8.

26. Rawji, K.S., Neumann, B., and Franklin, R.J.M. (2023). Glial aging and its impact on central nervous system myelin regeneration. Ann. N. Y. Acad. Sci. 1519, 34–45. 10.1111/nyas.14933.

27. Diniz, L.P., Araujo, A.P.B., Carvalho, C.F., Matias, I., de Sá Hayashide, L., Marques, M., Pessoa, B., Andrade, C.B.V., Vargas, G., Queiroz, D.D., et al. (2024). Accumulation of damaged mitochondria in aging astrocytes due to mitophagy dysfunction: Implications for susceptibility to mitochondrial stress. Biochim. Biophys. Acta - Mol. Basis Dis. 1870. 10.1016/j.bbadis.2024.167470.

28. Katayama, H., Kogure, T., Mizushima, N., Yoshimori, T., and Miyawaki, A. (2011). A sensitive and quantitative technique for detecting autophagic events based on lysosomal delivery. Chem. Biol. 18, 1042–1052. 10.1016/j.chembiol.2011.05.013.

29. Violot, S., Carpentier, P., Blanchoin, L., and Bourgeois, D. (2009). Reverse pH-Dependence of Chromophore Protonation Explains the Large Stokes Shift of the Red Fluorescent Protein mKeima. 10356–10357.

30. Henderson, J.N., Osborn, M.F., Koon, N., Gepshtein, R., Huppert, D., and Remington, S.J. (2009). Excited State Proton Transfer in the Red Fluorescent Protein mKeima. 13212–13213.

31. Kogure, T., Kawano, H., Abe, Y., and Miyawaki, A. (2023). Fluorescence imaging using a fluorescent protein with a large Stokes shift dKeima. 45, 223–226. 10.1016/j.ymeth.2008.06.009.

32. Kawano, H., Kogure, T., Abe, Y., Mizuno, H., and Miyawaki, A. (2008). Two-photon dual-color imaging using fluorescent proteins. 5, 373–374.

33. Fang, E.F., Hou, Y., Lautrup, S., Jensen, M.B., Yang, B., SenGupta, T., Caponio, D., Khezri, R., Demarest, T.G., Aman, Y., et al. (2019). NAD+ augmentation restores mitophagy and limits accelerated aging in Werner syndrome. Nat. Commun. 10. 10.1038/s41467-019-13172-8.

34. López-Otín, C., Blasco, M.A., Partridge, L., Serrano, M., and Kroemer, G. (2023). Hallmarks of aging: An expanding universe. Cell 186, 243–278. 10.1016/j.cell.2022.11.001.

35. Schmauck-Medina, T., Molière, A., Lautrup, S., Zhang, J., Chlopicki, S., Madsen, H.B., Cao, S., Soendenbroe, C., Mansell, E., Vestergaard, M.B., et al. (2022). New hallmarks of ageing: a 2022 Copenhagen ageing meeting summary. Aging (Albany. NY). 14, 6829–6839. 10.18632/aging.204248.

36. Jiménez-Loygorri, J.I., Villarejo-Zori, B., Viedma-Poyatos, Á., Zapata-Muñoz, J., Benítez-Fernández, R., Frutos-Lisón, M.D., Tomás-Barberán, F.A., Espín, J.C., Area-Gómez, E., Gomez-Duran, A., et al. (2024). Mitophagy curtails cytosolic mtDNA-dependent activation of cGAS/STING inflammation during aging. Nat. Commun. 15. 10.1038/s41467-024-45044-1.

37. Nieuwenhuis, B., Haenzi, B., Hilton, S., Carnicer-Lombarte, A., Hobo, B., Verhaagen, J., and Fawcett, J.W. (2021). Optimization of adeno-associated viral vector-mediated transduction of the corticospinal tract: comparison of four promoters. Gene Ther. 28, 56–74. 10.1038/s41434-020-0169-1.

38. Schmid, E.T., Pyo, J.H., and Walker, D.W. (2022). Neuronal induction of BNIP3-mediated mitophagy slows systemic aging in Drosophila. Nat. Aging 2, 494–507. 10.1038/s43587-022-00214-y.

39. Hjukse, J.B., Puebla, M.F.D.L., Vindedal, G.F., Sprengel, R., Jensen, V., Nagelhus, E.A., and Tang, W. (2023). Increased membrane Ca2+ permeability drives astrocytic Ca2+ dynamics during neuronal stimulation at excitatory synapses. Glia, 1–12. 10.1002/glia.24450.

40. Tang, W., Szokol, K., Jensen, V., Enger, R., Trivedi, C.A., Hvalby, Ø., Johannes Helm, P., Looger, L.L., Sprengel, R., and Nagelhus, E.A. (2015). Stimulation-evoked Ca2+ signals in astrocytic processes at hippocampal CA3-CA1 synapses of adult mice are modulated by glutamate and ATP. J. Neurosci. 35, 3016–3021. 10.1523/JNEUROSCI.3319-14.2015.

41. Fang, E.F., Kassahun, H., Croteau, D.L., Scheibye-Knudsen, M., Marosi, K., Lu, H., Shamanna, R.A., Kalyanasundaram, S., Bollineni, R.C., Wilson, M.A., et al. (2016). NAD+ Replenishment Improves Lifespan and Healthspan in Ataxia Telangiectasia Models via Mitophagy and DNA Repair. Cell Metab. 24, 566–581. 10.1016/j.cmet.2016.09.004.

42. Hou, Y., Wei, Y., Lautrup, S., Yang, B., Wang, Y., Cordonnier, S., Mattson, M.P., Croteau, D.L., and Bohr, V.A. (2021). NAD+ supplementation reduces neuroinflammation and cell senescence in a transgenic mouse model of Alzheimer’s disease via cGAS-STING. Proc. Natl. Acad. Sci. U. S. A. 118. 10.1073/pnas.2011226118.

43. Zhang, H., Ryu, D., Wu, Y., Gariani, K., Wang, X., Luan, P., Amico, D.D., Ropelle, E.R., Lutolf, M.P., Aebersold, R., et al. (2016). Supplementary Materials for enhances life span in mice. Science (80-.). 352, 1436–1443.

44. Marmolejo-Garza, A., Chatre, L., Croteau, D.L., Herron-Bedoya, A., Luu, M.D.A., Bernay, B., Pontin, J., Bohr, V.A., Boddeke, E., and Dolga, A.M. (2024). Nicotinamide riboside modulates the reactive species interactome, bioenergetic status and proteomic landscape in a brain-region-specific manner. Neurobiol. Dis. 200, 106645. 10.1016/j.nbd.2024.106645.

45. Meyer, T., Shimon, D., Youssef, S., Yankovitz, G., Tessler, A., Chernobylsky, T., Gaoni-Yogev, A., Perelroizen, R., Budick-Harmelin, N., Steinman, L., et al. (2022). NAD+ metabolism drives astrocyte proinflammatory reprogramming in central nervous system autoimmunity. Proc. Natl. Acad. Sci. U. S. A. 119, 1–12. 10.1073/pnas.2211310119.

46. Khawaja, R.R., Martín-Segura, A., Santiago-Fernández, O., Sereda, R., Lindenau, K., McCabe, M., Macho-González, A., Jafari, M., Scrivo, A., Gomez-Sintes, R., et al. (2025). Sex-specific and cell-type-specific changes in chaperone-mediated autophagy across tissues during aging. Nat. Aging 5, 8–10. 10.1038/s43587-024-00799-6.

47. Shevtsova, Z., Malik, J.M.I., Michel, U., Bähr, M., and Kügler, S. (2005). Promoters and serotypes: Targeting of adeno-associated virus vectors for gene transfer in the rat central nervous system in vitro and in vivo. Exp. Physiol. 90, 53–59. 10.1113/expphysiol.2004.028159.

48. Kirchhoff, F., and Tang, W. (2024). Analysis of Functional NMDA Receptors in Astrocytes. Methods Mol. Biol. 2799, 201–223. 10.1007/978-1-0716-3830-9_11.

49. Pnevmatikakis, E.A., and Giovannucci, A. (2017). NoRMCorre: An online algorithm for piecewise rigid motion correction of calcium imaging data. J. Neurosci. Methods 291, 83–94. 10.1016/j.jneumeth.2017.07.031.

50. Berg, S., Kutra, D., Kroeger, T., Straehle, C.N., Kausler, B.X., Haubold, C., Schiegg, M., Ales, J., Beier, T., Rudy, M., et al. (2019). Ilastik: Interactive Machine Learning for (Bio)Image Analysis. Nat. Methods 16, 1226–1232. 10.1038/s41592-019-0582-9.

51. Rao, S.B., Katoozi, S., Skauli, N., Froehner, S.C., Ottersen, O.P., Adams, M.E., and Amiry-Moghaddam, M. (2019). Targeted deletion of β1-syntrophin causes a loss of K ir 4.1 from Müller cell endfeet in mouse retina. Glia 67, 1138–1149. 10.1002/glia.23600.

52. Chakraborty, J., Caicci, F., Roy, M., and Ziviani, E. (2020). Investigating mitochondrial autophagy by routine transmission electron microscopy: Seeing is believing? Pharmacol. Res. 160, 105097. 10.1016/j.phrs.2020.105097.

53. Mizushima, N., Yoshimori, T., and Levine, B. (2010). Methods in Mammalian Autophagy Research. Cell 140, 313–326. 10.1016/j.cell.2010.01.028.

54. Jung, M., Choi, H., and Mun, J.Y. (2019). The autophagy research in electron microscopy. Appl. Microsc. 49, 1–7. 10.1186/s42649-019-0012-6.

