## Supplemental Information for "Two-photon *in vivo* imaging reveals cell type-specific mitophagy dynamic changes in mouse somatosensory cortex during aging"

**Figure S1.** Related to Figure 1.

**Video S1.** Two-photon mt-Keima imaging of somatosensory cortical neurons in early-aged mice, related to Figure 1.

**Video S2.** Two-photon mt-Keima imaging of somatosensory cortical neurons in old-aged mice, related to Figure 1.

**Video S3.** Two-photon mt-Keima imaging of somatosensory cortical astrocytes in early-aged mice, related to Figure 1.

**Video S4.** Two-photon mt-Keima imaging of somatosensory cortical astrocytes in old-aged mice, related to Figure 1.

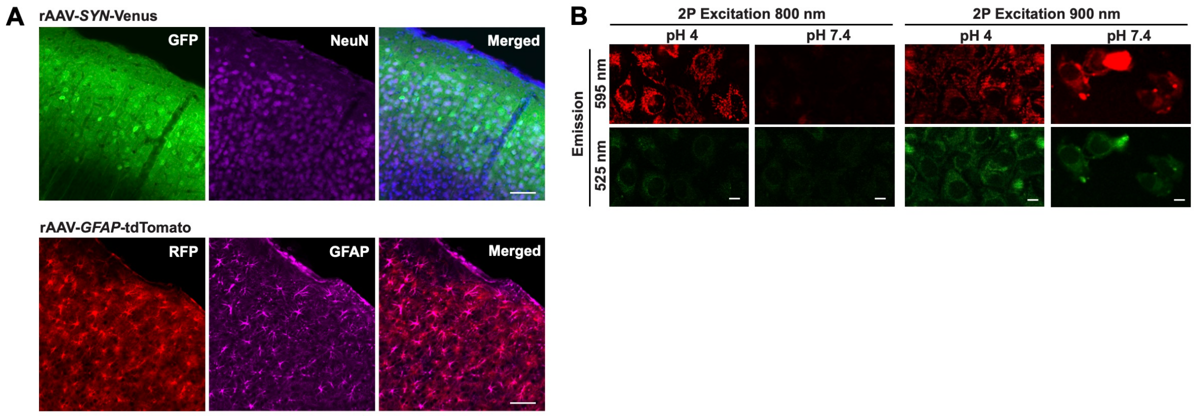


**Figure S1: (A) Immunohistochemistry showing cell-type specific labeling of neurons and astrocytes in mouse somatosensory cortex.** The human *SYN* and human *GFAP* promoters drive cell type-specific expression of Venus and tdTomato via rAAV transduction in neurons and astrocytes, respectively. The Venus and tdTomato positive cells are colocalized with neuronal marker NeuN and astrocytic marker GFAP, respectively. Scale bar, 50 µm. **(B)** Two-photon imaging of mt-Keima expressing Hela cell line under acidic (pH 4) and neutral (pH 7.4) environment with two-photon excitation wavelengths at 900 nm and 800 nm and recording in both green and red channels. Scale bar, 10 µm.
